# Open source modules for tracking animal behavior and closed-loop stimulation based on Open Ephys and Bonsai

**DOI:** 10.1101/340141

**Authors:** Alessio Paolo Buccino, Mikkel Elle Lepperød, Svenn-Arne Dragly, Philipp Häfliger, Marianne Fyhn, Torkel Hafting

## Abstract

**Objective:** A major goal in systems neuroscience is to determine the causal relationship between neural activity and behavior. To this end, methods that combine monitoring neural activity, behavioral tracking, and targeted manipulation of neurons in closed-loop are powerful tools. However, commercial systems that allow these types of experiments are usually expensive and rely on non-standardized data formats and proprietary software which may hinder user-modifications for specific needs. In order to promote reproducibility and data-sharing in science, transparent software and standardized data formats are an advantage. Here, we present an open source, low-cost, adaptable, and easy to set-up system for combined behavioral tracking, electrophysiology and closed-loop stimulation.

**Approach:** Based on the Open Ephys system (www.open-ephys.org) we developed multiple modules to include real-time tracking and behavior-based closed-loop stimulation. We describe the equipment and provide a step-by-step guide to set up the system. Combining the open source software Bonsai (`bonsai-rx.org`) for analyzing camera images in real time with the newly developed modules in Open Ephys, we acquire position information, visualize tracking, and perform tracking-based closed-loop stimulation experiments. To analyze the acquired data we provide an open source file reading package in Python.

**Main results:** The system robustly visualizes real-time tracking and reliably recovers tracking information recorded from a range of sampling frequencies (30-1000Hz). We combined electrophysiology with the newly-developed tracking modules in Open Ephys to record place cell and grid cell activity in the hippocampus and in the medial entorhinal cortex, respectively. Moreover, we present a case in which we used the system for closed-loop optogenetic stimulation of entorhinal grid cells.

**Significance:** Expanding the Open Ephys system to include animal tracking and behavior-based closed-loop stimulation extends the availability of high-quality, low-cost experimental setup within standardized data formats serving the neuroscience community.

## 1 Introduction

To understand how brain function arises from neural activity, it is helpful to utilize methods that both monitor groups of single-neurons at single-spike resolution and behavior [4]. While correlations between recorded neural activity and behavior or sensory stimuli reveal associative relations, targeted manipulations are necessary to identify cell-specific contributions and determine causal relations.

Combining electrophysiological recordings and positional tracking has led to seminal discoveries about the neural correlates of behaviors such as place cells [27], grid cells [14, 16], head-direction cells [32], border cells [35], and speed cells [20]. Moreover, behavioral tracking of freely moving rodents is also essential to reveal neural mechanisms underlying a range of behaviors, including aggression, social interactions, defensive behaviors, and different memory tasks [37, 21, 36].

The experimental setups necessary for such investigations require a combination of electrophysiology and tracking of animal behavior. Neural recordings are conducted with implanted electrodes recording extracellular fluctuations of electric potential. Tracking the animal’s position is usually done by mounting Light Emitting Diodes (LEDs) on the animal’s implant and tracking the position of LEDs with an optical camera.

A growing number of vendors provide solutions for experiments combining electrophysiology and animal tracking. We selected three representative vendors and asked for quotes for the instrumentation needed to perform the experiments. We included the neural acquisition system, one 32-channel headstage (+ cable), the tracking system (including the camera), and the software (excluding general purpose hardware/software - workstations - and installation costs). The quotes that we received from the companies ranged from 60,000 to 75,000 USD. The high cost of these commercially available systems is only one of the limitations. In fact, each of the proposed systems uses different proprietary data formats. This may compromise the reproducibility of results and limit data sharing [5]. However, the services provided with commercial systems include priority support, and often specialized engineering help included in the cost.

The advances in instrumentation and recording systems have expanded the possibility of neural interrogation by interacting by introducing real-time feedback control designed by the experimenter. Closed-loop experiments utilize readouts of neural activity or behavior to make real-time decisions about how/when/where to stimulate the neural tissue. Stimulation is usually performed by means of electrical pulses through the electrodes, or using optogenetics – a methodology that allows millisecond-scale optical control of neural activity in genetically identified cell-types during animal behavior [19]. Quite a few studies have performed spike-triggered electrical stimulation [12, 9, 18, 26, 29, 8], while fewer have used real-time tracking to electrically stimulate single cells [22, 10]. Regarding optical stimulation, most experiments in behaving animals to date can be categorized as open-loop. The goal of this work, therefore, is to implement open source modules to easily control closed-loop electrical or optical stimulation.

In the neuroscience community, there has lately been a call for a world-wide open source effort (http://www.opensourceforneuroscience.org/) to standardize methods and share data in order to make break-through advancements in such a complicated field [15]. Many open hardware and software solutions have been developed for various aspects of neuroscience, including acquisition systems for electrophysiology [33, 2], portable miniscopes [6, 24], software tools for real-time interface with external devices [25], and closed-loop systems [7, 33].

Among the open source systems for electrophysiology hardware and software, Open Ephys (www.open-ephys.org – [33]) has gained popularity in the past few years. The Open Ephys acquisition system can currently be interfaced with up to 512 recording channels that are sampled by an Opal Kelly XEM-6310 FPGA module, connected via a USB 3.0 port. The FPGA sends 10 ms epochs, and the GUI can process the data in buffers ranging from 3 ms to 42 ms. Closed-loop latencies are mainly due to USB communication, but the system is being constantly improved: as of May 2018, a version based on PCI express communication is under development, which would lower latencies to only hundreds of microseconds [33]. However, animal tracking has not been integrated directly with the Open Ephys system.

For tracking of animal behavior, there are also a number of open source software tools available, including GemVid [28], OpenControl [1], MouseMove [31], and others [17]. Most of these tools allow for tracking of animal behavior and complex analysis, but they lack either the capability of real-time interaction with the system, closed-loop experiments, or integration with electrophysiology. Thus, we propose an open source solution to perform experiments involving animal tracking in a closed-loop mode by extending on existing open source tools, namely Open Ephys and Bonsai (`bonsai-rx.org` – [25]).

Bonsai is an open source visual programming framework for processing data streams [25]. Bonsai alone can be used for streaming and visualization of electrophysiology and tracking data, including closed-loop stimulation. However, we chose to use Bonsai only to extract positional information from the camera, and rather expand the Open Ephys GUI because it includes a multitude of useful plugins for electrophysiology and is widely used by the electrophysiology community.

Our system represents a cheap (< 5,000USD), reproducible, and customizable alternative solution to commercially available systems. It also allows tracking-based closed-loop stimulation that can be used with electrical or optogenetic stimulation.

In the following sections, we present an overview of the system before describing the image processing performed with Bonsai and introducing the newly implemented Open Ephys plugins for tracking and closed-loop stimulation. Then, we show results on the performance of this arrangement, first in open-field acquisitions of behavioral experiments involving place and grid cells, and finally, on a setup with closed-loop tracking-based stimulation.

## 2 Materials and Methods

In this section we provide an overview of the system, a description of the Bonsai script, and a detailed presentation of the plugins for the Open Ephys GUI. The source code is published on GitHub (https://github.com/CINPLA/tracking-plugin and https://github.com/CINPLA/logic-gate-plugin), with a wiki page containing a detailed guide on how to set up the system (https://github.com/CINPLA/tracking-plugin/wiki). A collection of Open Ephys plugins developed by our group, including those presented in this paper, is available at https://github.com/CINPLA/open-ephys-plugins.

### 2.1 System overview

Figure 1 gives an overview of the proposed solution at the systems level. The animals are implanted with any neural probe with an Omnetics connector that interfaces with an Intan RHD2000 series chip (compatible with the Open Ephys acquisition board). Two Light Emitting Diodes (LEDs) - in our case a red and a green off-the-shelf LED - are mounted on the headstage and are tracked during the experiment to extract the animal’s position. By using two separate light sources instead of one we can extract the relative angle between them and determine the head direction and angular motion.

**Figure 1:**
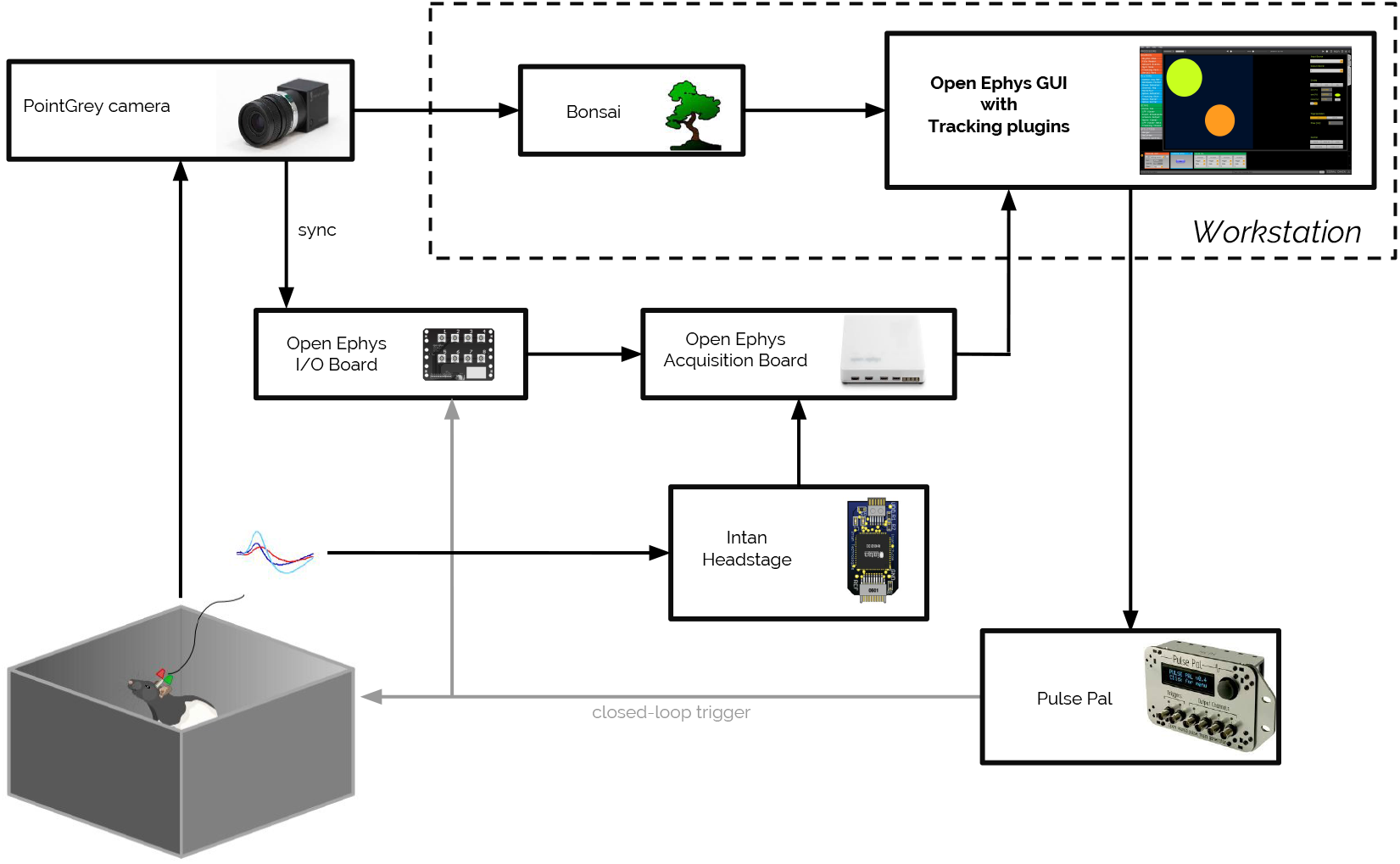
System level block diagram: the animal is equipped with recording electrodes and two LEDs. The electrophysiology data are recorded with the Open Ephys acquisition board (via the Intan headstage) and sent to the Open Ephys GUI. The tracking data are measured with a PointGrey camera and processed with Bonsai, which sends positions to the Open Ephys GUI. For closed-loop experiments, the Open Ephys GUI controls the Pulse Pal stimulator, whose pulses are also recorded using the Open Ephys I/O board. Synchronization between tracking and electrophysiology is performed by recording the camera shutter events using the Open Ephys I/O board.

The frames from the camera (we used a PointGrey Flea3 camera mounted in the ceiling pointing downwards to the box in which the rodents move around), are analyzed by Bonsai, in which an image processing pipeline is implemented to extract the position of the LEDs. The tracking data are sent to the Open Ephys GUI for saving and visualization, as well as to trigger tracking-based stimulation.

Electrophysiology data are measured by the Intan headstage (RHD series, or any other compatible with the Open Ephys acquisition board) and sent via Serial Peripheral Interface (SPI) cables to the acquisition system, which inputs the data stream to the Open Ephys GUI.

Synchronization between the electrophysiology and tracking systems is performed by recording Transistor-Transistor-Logic (TTL) events sent from the camera for every frame acquired, that is, every time the camera shutter closes (shutter TTL events), with the Open Ephys board.

The proposed system includes newly-developed *ad hoc* plugins for the Open Ephys GUI to record and visualize tracking data, and for closed-loop stimulation based on tracking. We suggest to use the

Pulse Pal stimulator to trigger external devices via TTL pulses, as it is easily interfaced to the Open Ephys system, but other devices could be used, such as Arduino.

Table 1 contains the components needed to set up the tracking-electrophysiology system, including company, vendor website, and cost. With the suggested system, an experiment combining electrophysiology and animal tracking can be set up with less than 5,000 USD (excluding the neural probe, but including a 32-channel headstage).

### 2.2 Image processing in Bonsai

Bonsai is a visual language designed for making software systems that require rich and rapid interaction with the external world [25]. In our setup we use it to grab frames from the camera, choose the region of interest, extract the x and y positions, and send them to the Open Ephys GUI via Open Sound Control (OSC) messages.

The image processing pipeline is shown in Figure 2 and it is performed by the `tracking.bonsai` script (in the *Bonsai* folder of https://github.com/CINPLA/tracking-plugin). Frames are streamed through the *FlyCapture* module and the *Crop* node allows users to resize the video so that it fits the area in which the rodents are roaming. Then, two custom-made color filter nodes (*Green* and *Red)* recognize green and red colors, respectively (colors can be easily changed by adjusting the parameters). The *Threshold* node sets a binary threshold to select the colored light source, and the *FindContours*, *Binary Region Analysis*, and *LargestBinaryBlob* nodes identify the blob position in the image. The centroids of red and green regions *(Source. Centroid)* are then zipped together with the size of the field view, that is, the width and height.

**Figure 2:**
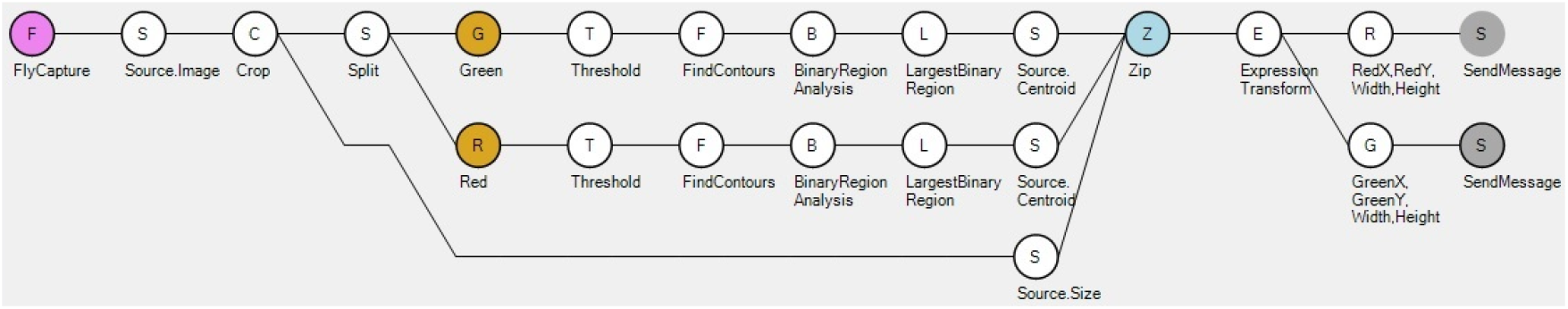
Bonsai image processing diagram. The images are acquired from the camera *(FlyCapture*) and the region of interest can be interactively selected *(Crop*). Green and red areas are then identified with color filters *(Green* and *Red*) and then thresholded *(Threshold* – thresholds can be adjusted by the user) to compute the centroid of the their region. The x and y positions are then normalized between 0 and 1 dividing them by the width and height of the region of interest, respectively, and finally transmitted using the Open Sound Control (OSC) protocol *(SendMessage*).

**Table 1:** System components: each table row shows the component’s name, the company (or project) providing the component, the vendor website, and the cost in USD. The entire system, including a 32-channel headstage and cable for electrophysiology, can be acquired for less than 5,000 USD.

| Component | Company | Vendor Site | Cost |
| --- | --- | --- | --- |
| *Camera* |  |  |  |
| PointGrey Camera Flea3 | Point Grey | www.ptgrey.com | 599.95 USD |
| *Headstage* |  |  |  |
| Intan Headstage RHD2132 | Intan | intantech.com | 895 USD |
| Intan SPI cable (1.8 m) | Intan | intantech.com | 295 USD |
| *Acquisition system* |  |  |  |
| Open Ephys Acquisition Board | Open Ephys | www.open-ephys.org | 2,500 USD |
| Open Ephys I/O Board | Open Ephys | www.open-ephys.org | 15 USD |
| *Stimulator* |  |  |  |
| Pulse Pal | Sanworks | www.sanworks.io | 595 USD |
| *Software* |  |  |  |
| Bonsai | Bonsai | bonsai-rx.org | 0 USD |
| Open Ephys GUI | Open Ephys | www.open-ephys.org | 0 USD |
| Total |  |  | 4,899.95 USD |

The following part of the pipeline deals with the OSC communication. First, the x and y positions are normalized *(ExpressionTransform*), that is, divided by the width and height of the cropped regions, respectively. Finally, they are packed into two OSC messages: one for the red LED and one for the green LED. Each message contains 4 float values: x, y, width and height.

We provide an extra Bonsai script (`osc.bonsai`), that sets up a RedPort (`port = 27020, address = “/red”, ip address` = “localhost”) and a GreenPort (`port = 27021, address` = “/green”, `ip address` = “localhost”) to speed up the system configuration. The user can change the ports, addresses and IP addresses in the *Properties* tab (currently, the Open Ephys tracking plugin can only receive OSC messages from “localhost”).

Although we provide Bonsai scripts for tracking, any tracking system is compatible with the Open Ephys plugin, as long as it sends OSC data in the format of 4 `float` values: x, y, width and height. The tracking data are then input to the Open Ephys GUI using the newly designed modules described in the next section.

### 2.3 Open Ephys GUI tracking plugin

We developed three modules publicly available at https://github.com/CINPLA/tracking-plugin/ (in the *Tracking* folder) to stream and save OSC tracking data into the GUI (`Tracking Port`), to visualize the path trajectories (`Tracking Visualizer`), and to trigger tracking-based closed-loop stimulation (`Tracking Stimulator`).

#### 2.3.1 Tracking Port

The `Tracking Port` is a *Source* of the Open Ephys signal chain, as it adds data into the GUI data buffers and saves the tracking signals. The module consists of an Editor only, displayed in Figure 3, which allows the user to add and delete *Tracking Sources* with the + and - buttons and to set the port, address, and color for each added source. The underlying processor robustly handles the streaming and circulation of the tracking data using two helper classes: the `Tracking Server` and the `Tracking Queue`.

**Figure 3:**
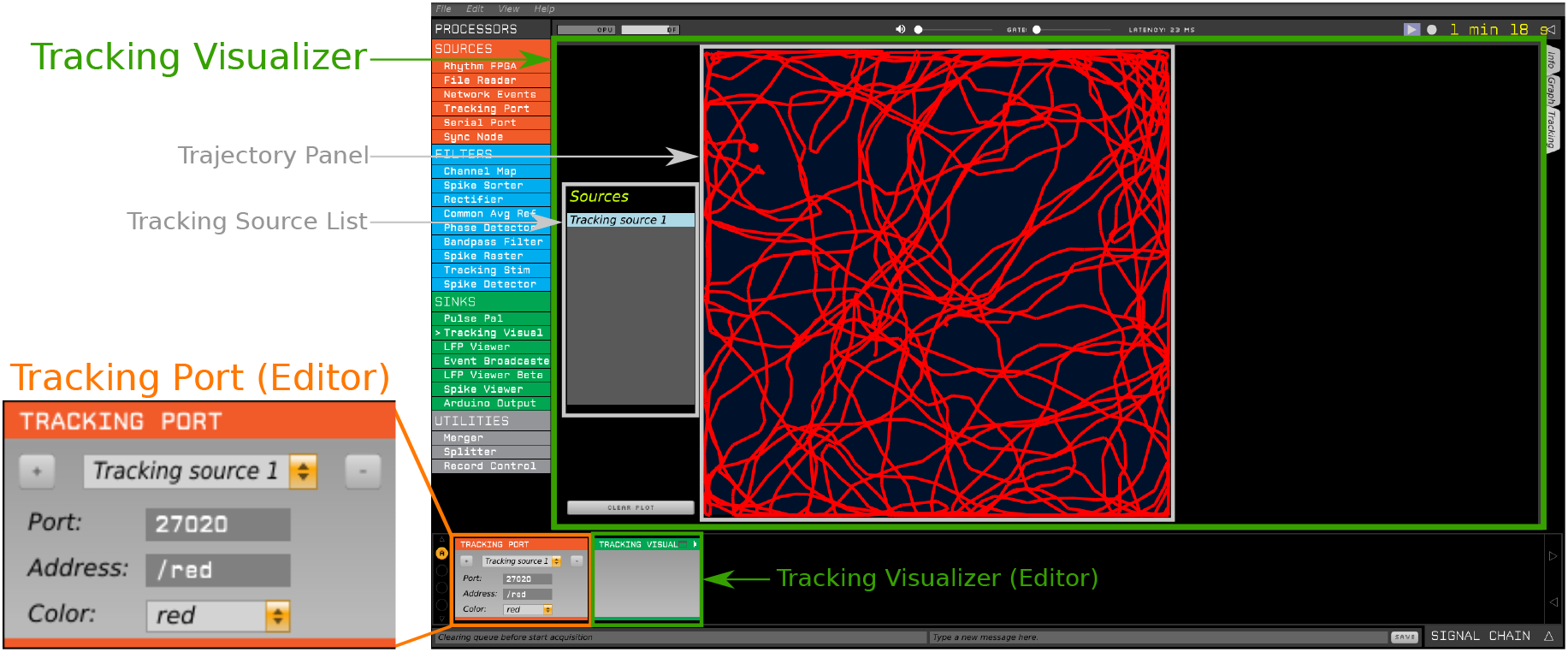
Open Ephys signal chain to visualize tracking data. The `Tracking Port` is a *Source* that allows the user to add and delete incoming sources (+ and -), to set the connection parameters for each tracking source (`Port` and `Address`) and to choose in which color the selected source will be displayed with the `Tracking Visualizer`. The `Tracking Visualizer` is a *Sink* module that permits visualization of the path trajectories from the sources set up in the `Tracking Port`. All the available sources are listed in the *Sources* list on the left and the user can select multiple sources at once. The path is represented in the trajectory panel. The CLEAR PLOT button can be used to erase the displayed trajectory (only for visualization purposes: the data are saved from the `Tracking Port` module).

For each new *Tracking Source*, a `Tracking Server` instance is created. The `Tracking Server` object takes care of the OSC communication and streams the data through a User Datagram Protocol (UDP) socket on a different thread, in order to make communication more robust. When the user changes the port or the address of a *Tracking Source*, the corresponding UDP socket is closed, the underlying thread is stopped, a new socket is opened, and a new thread is started. There is a maximum of 10 sources that can be streamed simultaneously on parallel threads.

The Open Ephys GUI processes data in buffers of around 20 ms at each processing cycle [33], which corresponds to 50 Hz. For some tracking systems this frequency could be too low, hence we implemented a queuing system that temporarily stores all received data and processes them in short intervals. Each *Tracking Source* has a `Tracking Queue` object that serves this purpose: when a new packet is received from the OSC data link, the tracking data and the timestamp at the time of reception are pushed in the queue, so that more than one message can be pushed for each source in the interval within processing cycles. When the `Tracking Port` processes the data (the `process()` function is called), the messages in the queue are popped and sent as binary events with the timestamp recorded at reception.

The combination of the `Tracking Server` and the `Tracking Queue` make data streaming very robust and reliable. In Section 3.1 we show how the system is able the correctly recover 10 different tracking sources sent at different frequencies between 30 and 300 Hz, as well as 10 high-speed 1 kHz sources.

#### 2.3.2 Tracking Visualizer

The `Tracking Visualizer` is a *Sink* module which is placed at the end of the signal chain to visualize the trajectories streamed by the `Tracking Port`. It is also a *Visualizer* module which can be docked in the GUI or opened as a separate window.

The *Visualizer* panel is shown in Figure 3: on the left side, a list of available sources is automatically updated and each source can be toggled for visualization (the maximum number of sources that can be visualized at the same time is 10). The tracking sources are plotted on the dark blue panel, whose proportions are adjusted depending on the width and height of the sources (for visualization purposes, the width and height are taken from the first available source only, but actual width and height are saved from the `Tracking Port` processor). The `CLEAR PLOT` button on the bottom left clears the displayed trajectories.

If some tracking information is missing, for example if the animal somehow covers the LEDs or the LEDs stop being visible, the resulting `NaN` (Not a Number) tracking data are skipped from visualization, to ensure a smooth path trajectory.

#### 2.3.3 Tracking Stimulator

The `Tracking Stimulator` is a *Filter* module and it allows the user to perform tracking-based closed-loop experiments. It was originally designed to stimulate place cells and grid cells inside or outside their fields with electrical or optogenetics stimulation.

It has a *Visualizer* panel, shown in Figure 4, that allows the user to manually draw, drag, resize (double click), copy (`ctrl+c`), paste (`ctrl+v`), and delete (`del`) circles, or input circles’ information (x_position, y_position and radius - from 0 to 1) using the editable labels on the right. Each circle can be inactivated using the `ON` toggle button.

**Figure 4:**
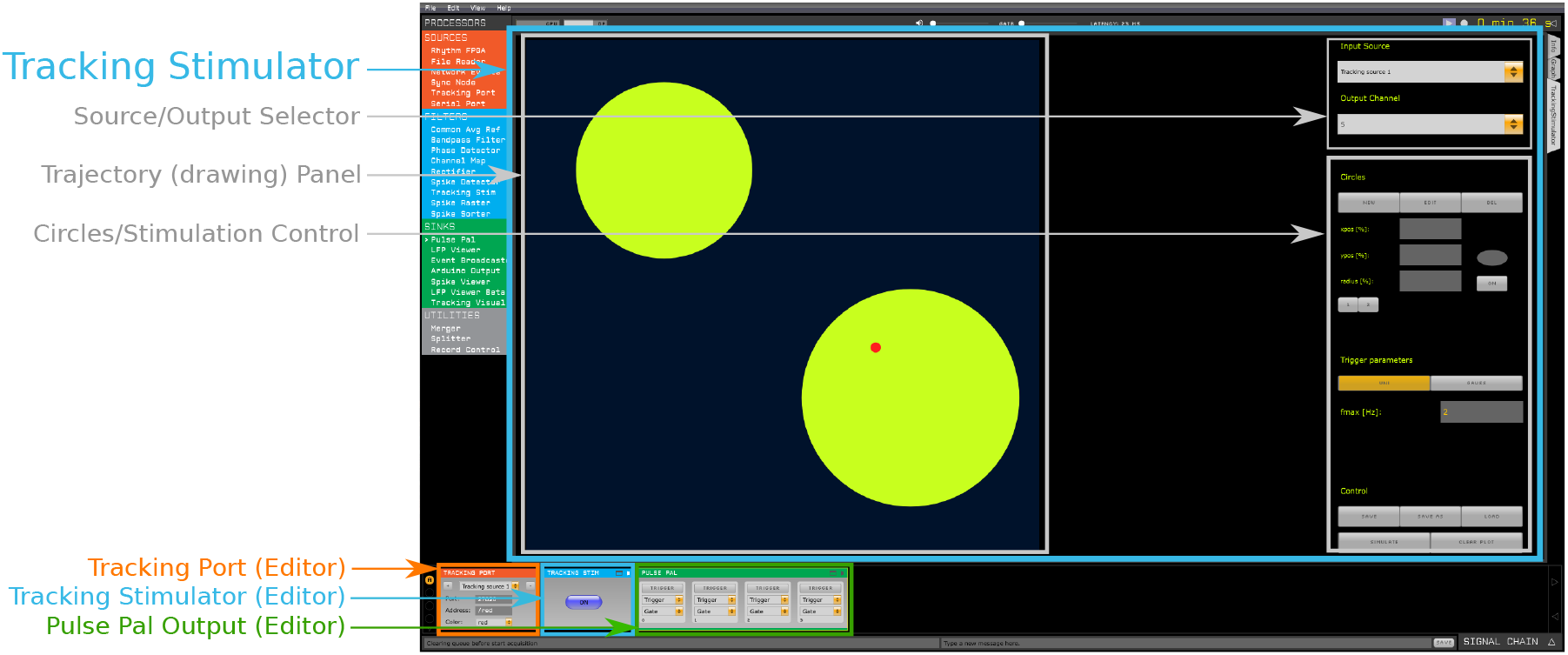
Open Ephys signal chain for closed-loop stimulation. The `Tracking Port` inputs the tracking data. The `Tracking Stimulator` is a *Filter* module used to perform closed-loop tracking-based stimulation. The user can draw circles in the trajectory (drawing) panel, and when the position of the selected source *(Input Source*) is within the area(s), a TTL event is sent on the selected *Output channel*. In this example, the TTL event is picked up by the `Pulse Pal Output`, that in turn triggers external hardware stimulation.

On the top right, the user can decide which source to track among the available *Tracking Sources* – `Input source` drop-down list – and which TTL output channel to trigger – `Output channel` dropdown list – when the chosen *Tracking Source* is within the active circles.

Stimulation is triggered only when the toggle button in the editor (bottom left) is set to `ON`. When within the selected circles, the tracking cue becomes red and TTL events are generated on the selected `Output channel`. There are two stimulation modes, uniform and gaussian, controlled by the buttons `UNI - GAUSS`:

1. UNIFORM: a TTL train with a constant frequency `fmax` defined by the user is generated when the position is within selected regions. In this mode, the colors of the circles are uniformly orange/yellow.
2. GAUSSIAN: the frequency of the TTL train is gaussian modulated. When the position is in the center of each circle, the frequency is `fmax`, when it is on the border of a circle the frequency is `sd * fmax`. In this mode, the colors of the circles are graded, darker in the center and lighter on the borders.

The uniform stimulation generates a train of TTL events with fixed stimulation period and it can be used, for example, to control a laser for optogenetics, in which a set of pulses with a certain frequency is usually used. The gaussian stimulation, instead, produces a stochastic TTL train, as stimulation is triggered *randomly* with a probability following a gaussian distribution. This kind of stimulation can be used to electrically stimulate a place or grid cell with variability similar to that experienced in physiological settings.

### 2.4 Logic Gate plugin

The `Tracking Stimulator` module alone allows the user to select circular regions and trigger stimulation within the selected areas with uniform or gaussian trains. However, some experimental protocols require a more advanced control flow to trigger stimulation. In order to improve customization of experiments and possibly combine tracking and electrophysiology data, we have implemented the `Logic Gate` plugin (https://github.com/CINPLA/logic-gate-plugin), shown in Figure 5.

**Figure 5:**
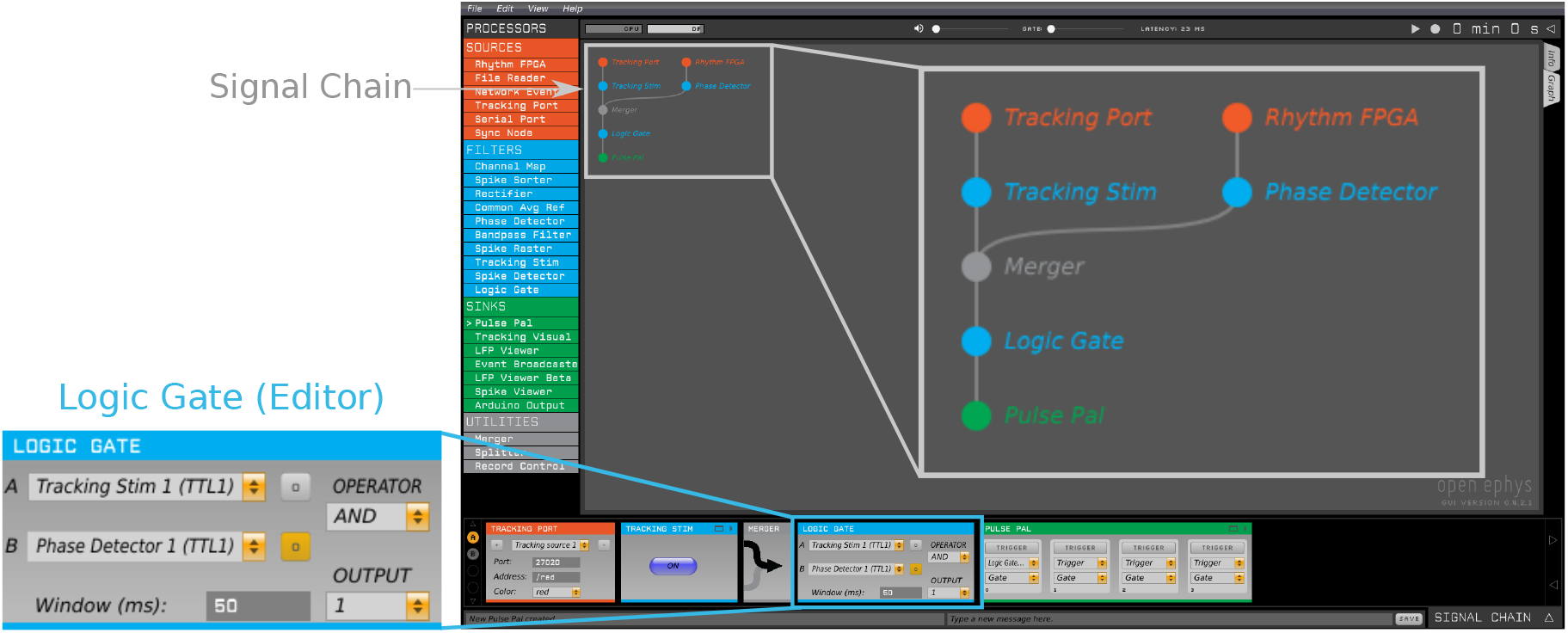
Open Ephys signal chain for closed-loop stimulation based on combined tracking and electrophysiology. The `Logic Gate` is a *Filter* module that allows the user to combine multiple TTL inputs and apply logical operators (and delay). In this example, stimulation is provided only when the animal is in the region selected with the `Tracking Stimulator` only after certain phase is detected on the neural signals by the `Phase Detector` (because signal `B`, corresponding to the `Phase Detector`, acts as gate).

The `Logic Gate` is a *Filter* module that allows the user to combine two TTL inputs with logical operators. To combine more than two inputs, multiple logic gates can be connected in series. The drop-down lists on the left side (`A` and `B`) allow users to select input signals. The available logical operators are `AND, OR, XOR`, and `DELAY`. The operator and output channel can be selected from the `OPERATOR` and `OUPUT` drop-down lists, respectively.

A TTL event is sent on the output channel if the events received on the input channels satisfy the selected logical operator within the specified time `Window`. The time window begins whenever an event is received on the channel defined as the gate, which is toggled by the buttons marked “ o” next to the input channels. If neither or both gate buttons are selected, both TTL inputs are used as gates and a new time window begins when an event is received on either channel. For example, if `B` serves as gate, `AND` is the logical operator, `Window` is set to 50ms, and `OUTPUT` is channel 1 (as in Figure 5), when a `B` event is received, only if an `A` signal is received within 50 ms an output TTL event will be sent on channel 1, but not vice versa (when `A` precedes `B`).

When `AND` is selected, as soon as the condition is satisfied within the `Window` a TTL output is generated; when `OR` or `XOR` are selected, the TTL event is sent at the end of the `Window` time if the logical condition is met (to propagate the signals instantaneously, the user can set `Window` to 0). When `DELAY` is selected, the `A` TTL is propagated to the output with a delay of `Window` ms.

In Figure 5, we show a simple example on how to use the `Logic Gate` to performed closed-loop stimulation depending on the animal position and the neural signal phase. The `Tracking Stimulator` and the `Phase Detector` outputs are merged to the `Logic Gate`, and the `Phase Detector` input (`B`) serves as gate. When the neural signal is at a certain phase, e.g. the *peak* phase, input `B` is received. For the following 50 ms, if the animal is in a selected region of the `Tracking Stimulator` and `A` inputs are received, then the `Logic Gate` will output events and, in this example, trigger stimulation through the `Pulse Pal Output`.

The combination of the Tracking and the `Logic Gate` plugins extends the possibilities for closed-loop paradigms within the Open Ephys environment.

### 2.5 Parsing the output files

In order to ease the set up of the proposed system and quickly access the recorded electropysiology and tracking data, we provide the `pyopenephys` Python package (https://github.com/CINPLA/py-open-ephys, which can also be accessed as a submodule from the main Tracking repository – https://github.com/CINPLA/tracking-plugin – in the *py-open-ephys* folder) to parse the output files into a Python environment with a few lines of code:

~~~
import pyopenephys
oe_file = pyopenephys.File(“path-to-recording-folder”)
experiment1 = oe_file.experiments[0]
recording1 = experiment1.recordings[0]
analog_signals = recording1.analog_signals
events_data = recording1.events
tracking_data = recording1.tracking
~~~

Once the data are loaded, it is straightforward to plot, for example, the trajectory of *Tracking Source 1*:

~~~
import matplotlib.pylab as plt
source_1 = tracking_data[0]
plt.plot(source_1.x, source_1.y)
~~~

The `pyopenephys` package also allows the user to easily export the analog signals, tracking data, and events of a recording to MATLAB format:

~~~
recording1.export_matlab(‘open-ephys.mat’)
~~~

The output file (`open-ephys.mat`) can be then loaded in the MATLAB environment and the Open Ephys tracking data can be accessed and plotted as follows:

~~~
*% Load data (times, duration, analog, tracking, and events variables)*
load(‘open-ephys.mat’);
*% Plot tracking source 1*
source_1 = squeeze(tracking(1, :, :));
x = source_1(1, :);
y = source_1(2, :);
plot(x, y);
~~~

## 3 Results

### 3.1 Reliability benchmark of tracking data transmission

First, we tested the reliability of the OSC transmission from the tracking system to the Open Ephys GUI. While positional tracking of animals is often limited to one or two tracking inputs, parallel processing of more channels opens for more detailed tracking information or simultaneous tracking of several animals. Therefore, we simulated a set of 10 random walks in Python and used the python-osc package to send them to the Open Ephys GUI at different frequencies equally spaced between 30 Hz to 300 Hz (denoted as *low frequency*) and another set of 10 sources at 1kHz (denoted as *high frequency*) (duration=60s). We recorded the received tracking data from the GUI and we recovered them using the pyopenephys script. Then, we compared the tracking data saved from the Python script (before the OSC link) and from the Open Ephys GUI (after the OSC link). The tests were run on an Ubuntu 16.04, Intel i7-6600U, 2.60GHz, 16 GB RAM machine under normal use with other ethernet traffic.

The data were almost perfectly recovered as all sent OSC packets were received and saved by the Open Ephys GUI (99 000 OSC messages sent/received for *low frequency* sources, 600 000 for *high frequency* sources).

In order to control the accuracy of timestamps recorded by Open Ephys we calculated the intervals between consecutive timestamps *(differential timestamps*) and compared the differential timestamps between the Python script and Open Ephys. In the histograms in Figure 6A we show the pairwise error between the sent and received differential timestamps for the 10 different sources at *low frequency*. The 99% confidence interval is [–2.6, 2.6] μs. Almost all of these errors are lower then 0.25 ms. Out of 98 990 errors, only 2 were more than 10ms (0.002%), 4 more than 5 ms (0.004%), 425 above 1ms (0.43%), and 1009 more than 500μs (1.02%). We found no significant differences among frequencies (for each pair of frequencies we used the Mann Whithney U test, finding effect sizes<10^−4^ – *Cohen’s d* coefficients – in all cases).

**Figure 6:**
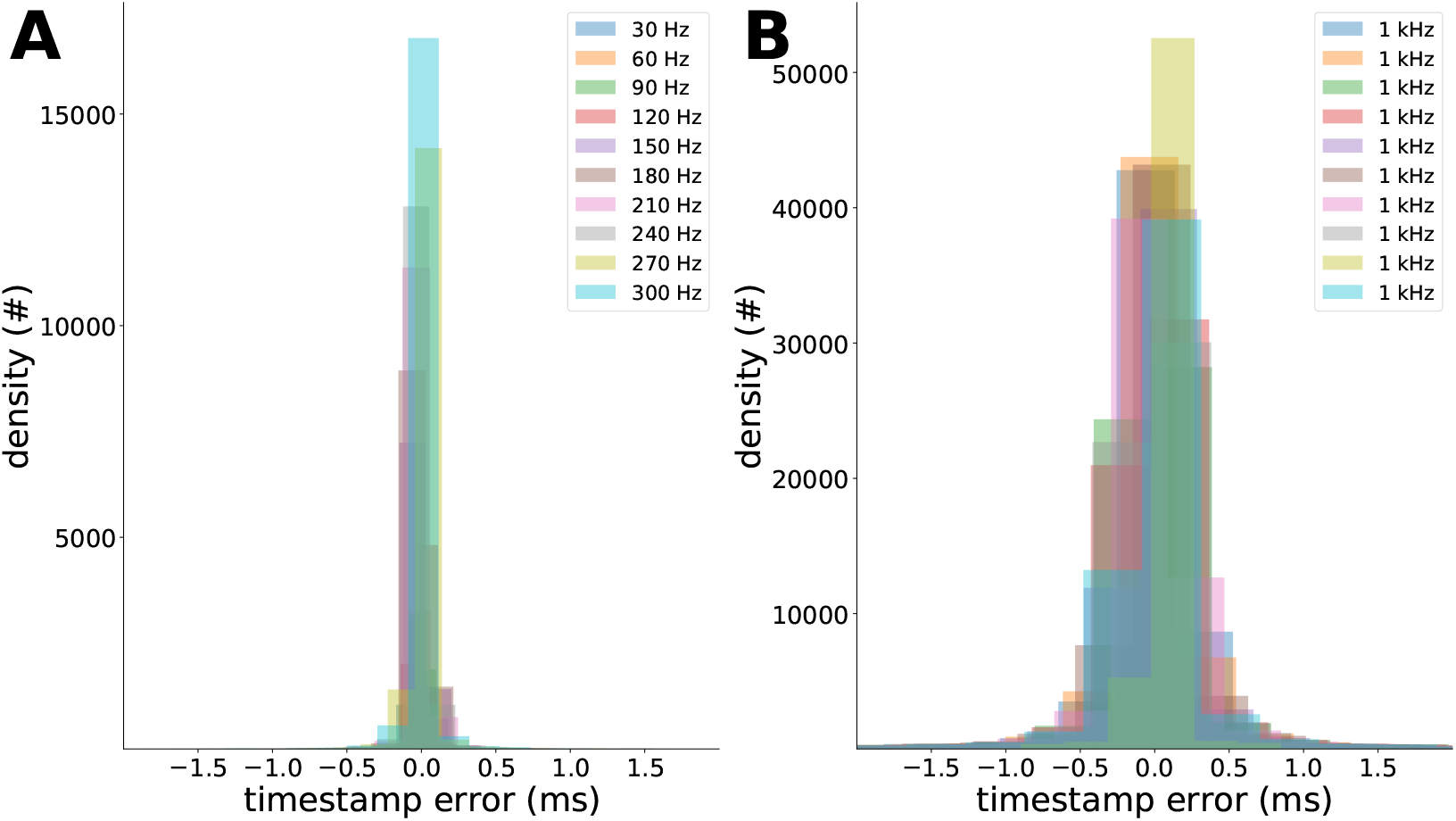
Histograms of the differential timestamp errors (for clarity we discarded errors above 10 ms – 2 observations for the *low frequency* errors and 224 for the *high frequency* ones). (A) For *low frequency* sources (10 sources from 30 Hz to 300 Hz), the 99% confidence interval is [–2.6, 2.6] μs and there is no worsening in performance as the frequency increases. (B) For *high frequency* sources (10 sources at 1 kHz), the error histograms are wider, with a 99 % confidence interval of [–5.22, 5.22] μs.

For the *high frequency* sources, in Figure 6B, the histograms are slightly wider. In this case the 99 % confidence interval is [–5.22, 5.22] μs. Only 224 out of 599 990 errors were more than 10 ms (0.037%), 3 309 more than 5ms (0.55%), 27 363 above 1ms (4.56%), and 54 200 more than 500μs (9.03%). In order to investigate whether errors accumulate over time, we compared the distributions of the first half of the received errors with the second half (299 995 observations each), but no significant difference was found (Mann Whitney U test).

These results show that the `Tracking Port` module is able to robustly handle multiple sources, even at high frequencies thanks to multi-threading (each *Tracking Source* is received in its own thread) and the `Tracking Queue`. The robust performance at high frequencies makes the system also suitable for tracking behaviors that require high-speed cameras, such as whisking or saccading [34].

### 3.2 Place and grid cells

We used the system in open-field experiments while recording neural activity in hippocampus and medial entorhinal cortex. Two adult Long Evans male rats were implanted with wired tetrodes (17 *μm* diameter) with impedance around 200 *kΩ* mounted on Axona (Axona Ltd (UK) – http://www.axona.com/) micro-drives. All experiments were approved by the Norwegian Animal Research Committee (FDU) before initiation. For more details about the experimental procedures, refer to *Lensjø et al*. [23]. A custom-built connector was used to interface the Axona to the Omnetics connector that connects to the current system (Figure 1). Electrophysiology data were processed offline with a 300-3000Hz bandpass filter (Butterworth - 3rd order), spike-sorted using KlustaKwik, and manually inspected using Phy [30]. Firing maps were then computed by splitting the arena in equal sized bins with 1 cm bin-size, dividing the number of spikes by the time spent by the rat in each bin and convolving with a two dimensional Gaussian kernel [14, 16].

Figure 7 displays the running path of the rat, spiking activity, and firing map of a place cell in the hippocampus CA1 area (A-B) and of a grid cell in medial entorhinal cortex (C-D). The duration of the experiments was approximately 10 minutes.

**Figure 7:**
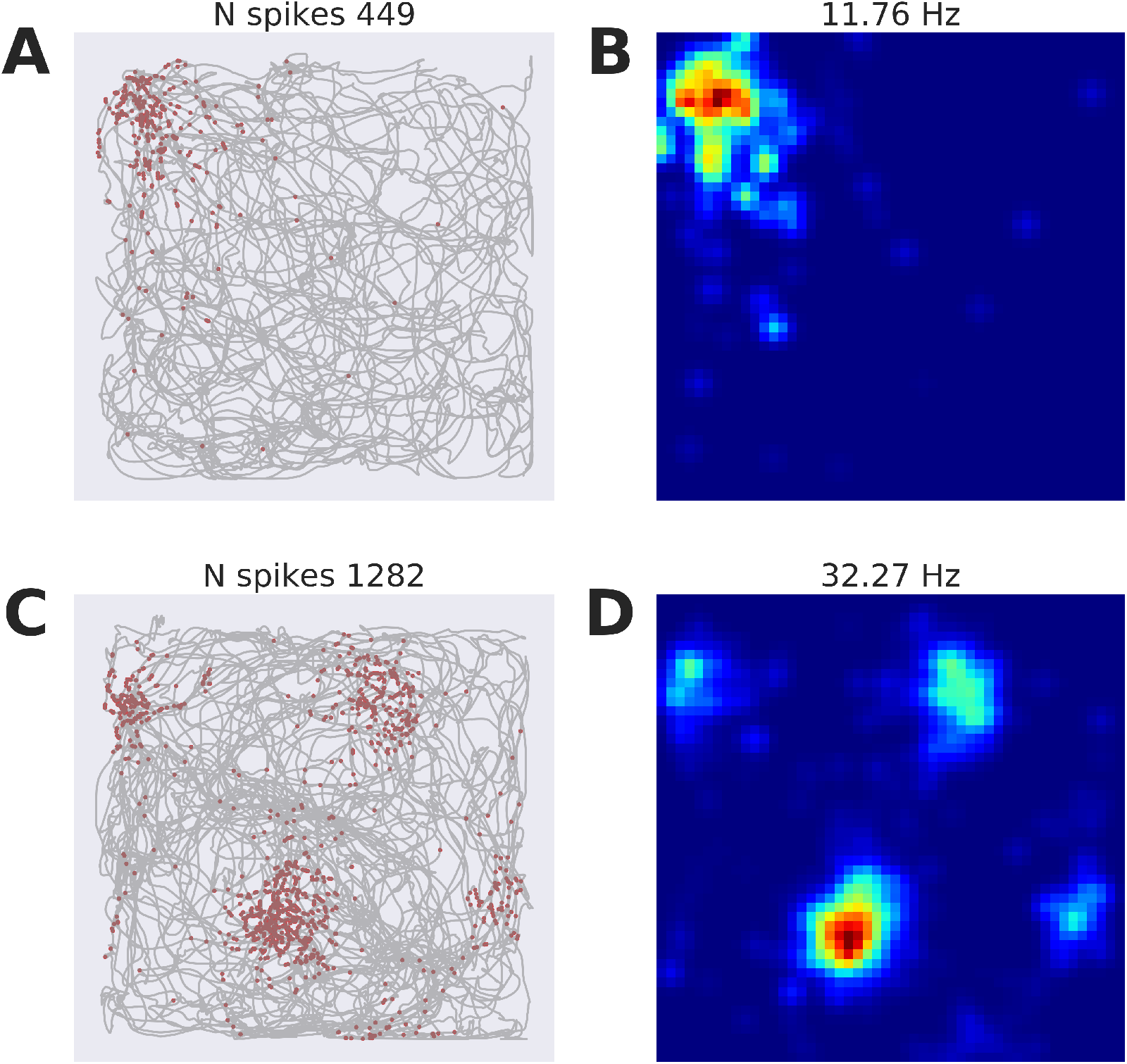
Place and grid cells recorded with the described system. (A-B) Place cell recorded from rat hippocampus in CA1 area. (A) Trajectory of the animal (grey line) with spikes superimposed as red dots on the path. (B) Color-coded firing rate map of the same single unit (blue, no firing; red, peak firing, maximum firing rate=11.76Hz). (C-D) Grid cell recorded from rat medial entorhinal cortex. (C) Trajectory of the rat with spikes superimposed. (D) Firing rate map of the unit in (C) that shows firing fields of the grid cell (maximum firing rate=32.27Hz).

### 3.3 Tracking-based closed-loop stimulation

Finally, we tested the tracking-based closed-loop stimulation. We simulated one random walk and sent it as an OSC message to the Open Ephys GUI. `The Tracking` Port module received the OSC message and sent the tracking data to the `Tracking Stimulator` module, in which we built a place field-like stimulation area. The `Tracking Stimulator` was then interfaced with the `Pulse Pal Output` module to send TTL triggers. The Pulse Pal stimulator was physically connected to the Open Ephys I/O board, so that the stimulation TTL events were also recorded from the acquisition system (Figure 1 – *closed-loop trigger*). Combining the simulated path trajectories with the TTL events sent by the Pulse Pal and recorded by the Open Ephys system, we constructed *stimulation trigger maps* – as explained in the previous section, but using TTL triggers instead of spikes – for the uniform and gaussian stimulation modes.

Figure 8A-B show the path trajectory with the stimulation TTL occurrences and the *stimulation trigger map* for the place field-like area using uniform stimulation (`fmax`=20Hz). In Figure 8C-D we used gaussian stimulation with `fmax`=20Hz and `sd`=0.1 % (2Hz at the borders). The recorded TTL signals, as expected, are almost perfectly uniformly distributed when uniform stimulation is applied (the lower rate on the borders is due to smoothing in computing the firing map) and gaussian-distributed when gaussian stimulation was selected. The duration of recording during the uniform stimulation was 1288 s, and for the gaussian stimulation it was 1741s.

**Figure 8:**
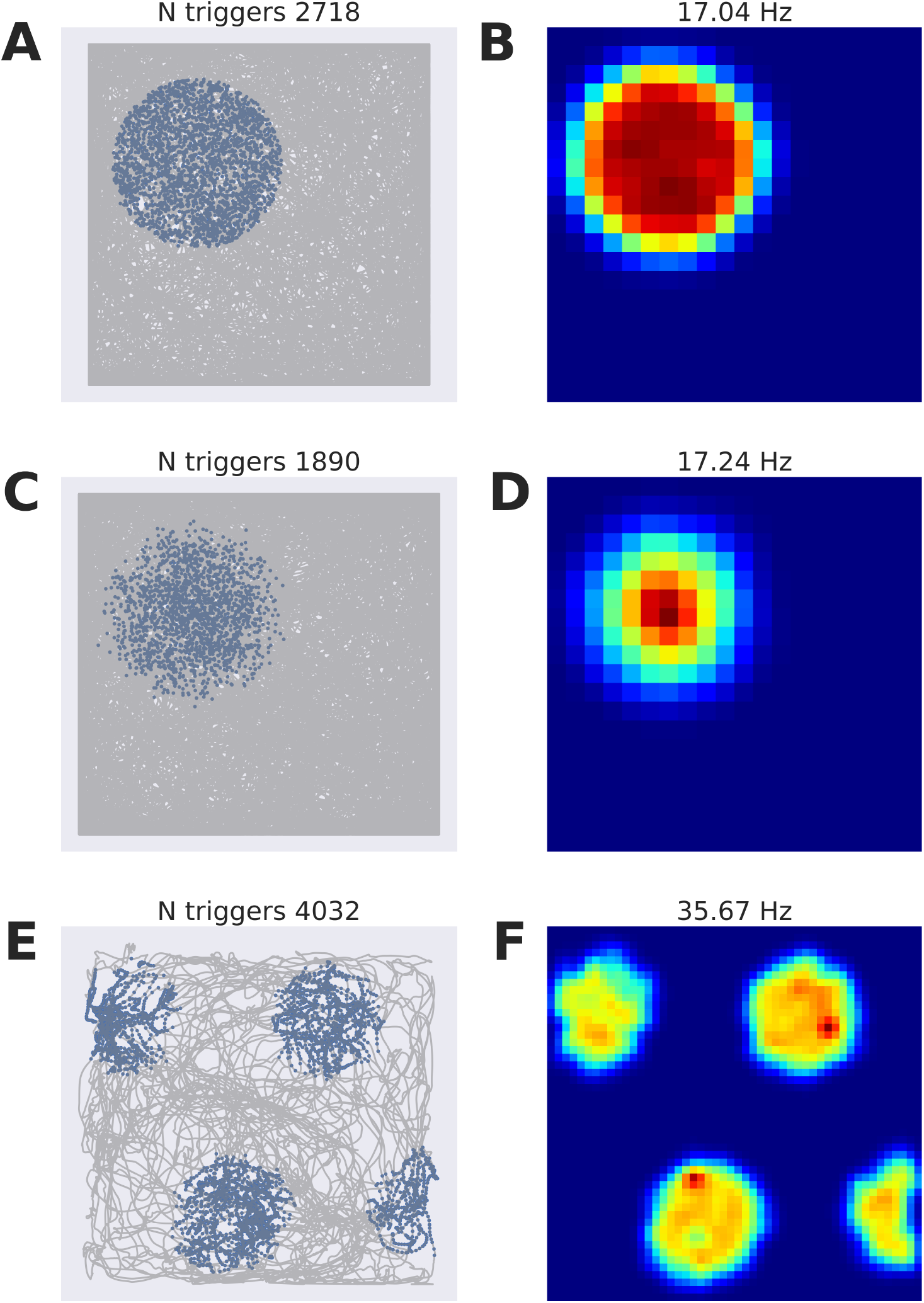
Closed-loop stimulation. Animal running paths in an open field were simulated with random walks and tracking information was sent as OSC messages to the Open Ephys GUI. A place field-like stimulation area was set in the `Tracking Stimulator` module and connected to the Pulse Pal. (AB) Uniform stimulation. (A) Path trajectory – grey line – and stimulation pulse occurrences – blue dots. (B) *Stimulation trigger map* showing an almost perfectly uniform distribution of the stimulation pulses (maximum stimulation rate=17.04Hz). (C-D) Gaussian stimulation. (C) Path trajectory and stimulation pulse occurrences. (D) *Stimulation trigger map* that shows a gaussian distribution of the stimulation pulses (maximum stimulation rate=17.24Hz). (E-F) Experimental grid cell stimulation. The grid cell depicted in Figure 7C-D was targeted with a grid field-like uniform stimulation pattern. (E) Animal path trajectory and stimulation pulse occurrences. (F) *Stimulation trigger map* that shows uniform grid pattern (maximum stimulation rate=|5.67 Hz)

We also used the uniform closed-loop stimulation to trigger blue laser pulses for selective activation of grid cells in medial enthorinal cortex. The grid cell shown in Figure 7C-D was used as target for stimulation. In Figure 8E-F we show the trajectory and *stimulation trigger map* of the laser pulses. The grid cell is stimulated in correspondence with its spatial firing fields (Figures 7D and 8F), but the stimulation fields are more uniform, as expected from the uniform simulation mode.

## 4 Discussions

In this work we presented an open source solution which integrates tracking of animal position using Bonsai [25] with electrophysiological recording and closed-loop stimulation using Open Ephys [33]. We developed plugins for the Open Ephys GUI which display the tracking of the animal and has the capability of performing closed-loop stimulation based on the animal’s position. We showed that the system is capable of handling multiple sources of tracking information at different frequencies and presented experimental data of an hippocampal place cell and a grid cell from the medial entorhinal cortex. We tested the closed-loop tracking based stimulation and showed that the extracted *stimulation trigger maps* are precise and accurate.

Closed-loop experiments involving spatial navigation are powerful tools to investigate memory formation and neuroplasticity. For example, *de Lavilléon et al*. [8] showed that using the spontaneous activity of a hippocampal place cell during sleep to trigger rewarding stimulations of the medial forebrain bundle (MFB) was able to induce a *place preference* towards the stimulated sites. Our system is not developed for spike-triggered stimulation, but can instead be used to stimulate the MFB during active behavior depending on the animal’s position. The approach to closed-loop stimulation that we presented here could represent an alternative to spike-triggered stimulation [12, 9, 18, 26, 29, 8].

Spike-triggered approaches require a fast online spike sorting and stimulation system, and works on the principle of inducing neuroplasticity using spike-timing dependent plasticity (STDP) protocols between a pair of coupled neurons. Instead, our closed-loop tracking-based stimulation is based on a behavior-driven stimulation, the principle of which is to induce neuroplasticity throughout the entire neural pathway, from behavior to neurons. Recently, for example, *Diamantaki et al*. [10] demonstrated the fast remapping of place cells with position-based closed-loop juxtacellular stimulation. In light of this, constraints on stimulation latency, which has led to the development of short-latency closed loop systems [7], have become more relaxed. The purpose of behavior-based stimulation, in fact, is to induce an average neural response and because the frequency of behavioral states (such as foraging, running, grooming) is much lower than the STDP window (10-50ms [18, 11]). Moreover, the implemented plugins allow to combine behavior and electrophysiology for closed-loop stimulation. One interesting possibility, given the importance of local field potential (LFP) oscillations for memory processing [3], is to use real-time position information and LFP phase to trigger stimulation (as demonstrated in Section 2.4). Such an approach could shed light on the role of precise timing in spatial navigation and memory formation.

The presented system represents an alternative to commercially available systems for animal tracking and neural recording. It comes at a cost of less than 10 % of the relevant commercial options, with an instrumentation cost just below 5,000 USD. The developed plugins are easy to use and the provided Python parser makes it very straightforward to access the raw data with a few lines of code (Section 2.5). Moreover, the system is fully open source with the source code available on GitHub (https://github.com/CINPLA/tracking-plugin and https://github.com/CINPLA/logic-gate-plugin), contributing to the commitment of the neuroscience community to provide open source data, instrumentation, and analysis tools [15]. While commercial solutions offer an integrated package, users of this system have to assemble and install hardware and software by themselves. A major caveat with open source solutions, in fact, is the support and maintenance of the systems. However, we chose to use Open Ephys and Bonsai because they represent excellent technical solutions, they are backed by renowned institutions, and they provide support from both a core group of developers and a growing community.

In the presented approach, we chose a specific vendor for the camera (PointGrey) and we picked a specific software for image processing (Bonsai). The choice of the camera was mainly dictated by the compatibility with Bonsai software and for its capability of sending precise TTL triggers every time the shutter closes (the TTL triggers are used for precise synchronization of tracking and electrophysiology data). Any other USB camera could be easily interfaced with Bonsai (by changing the video source in the image processing pipeline in Figure 2) and some of them would probably further cut the cost of the system, but this saving would come at the cost of lower resolution and frame rate, in addition to less precision in synchronization.

The choice of Bonsai was due to its partnership in the Open Ephys project and its proven versatility as a system for many different tracking scenarios, including 3D tracking of animal behavior. However, the communication between Bonsai and the Open Ephys GUI is achieved through the Open Sound Control (OSC) protocol and the Tracking plugin would work with any tracking data received in the specified format, such as the simulated data that we used in Sections 3.1 and 3.3. Moreover, we use and provide a Bonsai script that assumes that red and a green LEDs are mounted on the animal’s headstage. The Bonsai program could also be modified to extract the position by other means, such as a dark region on a light background, as is common in watermaze experiments [13]. Although Bonsai alone would have had the capability to interface with electrophysiology and tracking data, we decided to implement new Open Ephys modules and use Bonsai only to extract positional information from the video. This choice was made because the Open Ephys GUI is tailored for electrophysiology acquisition and a variety of plugins are already available for neural data analysis (`Spike Detector, Phase Detector, Bandpass Filter, Spike Sorting`, etc.) and visualization (`Spike Viewer` and `LFP Viewer`).

The current setup also presents some limitations. First, Bonsai is currently only supported on Windows machines, although the developers are planning to port it to Linux and macOS [25]. Another possible limitation regards electrophysiology and tracking data synchronization. As described above, we use TTL events from the camera shutter to realign the tracking data timestamps. However, in the current configuration, the camera in use sends a continuous flow of TTL shutter events as soon as it is switched on. Therefore, if the recording session starts (and Bonsai processing has started already) between the camera shutter time and the time Bonsai sends the OSC message, the Open Ephys system may receive an initial OSC tracking message *before* a TTL shutter event, due to image processing delay. In post processing, this eventuality can be corrected for by discarding OSC tracking messages received before TTL shutter events, and then pairing each TTL shutter event to the following received OSC tracking message. Ideally, this small problem would be avoided if the TTL flow of the camera could be started when Bonsai is started: in this scenario the user would start the Open Ephys recording and then the Bonsai pipeline, so that the first TTL event would correspond to the first OSC tracking message from Bonsai.

Another caveat is the use of UDP transmission for data communication, where the receiver does not send an *acknowledge* message to the sender and lost packages are not resent. Some messages could therefore be dropped. However, UDP is suitable for real-time applications due to its higher throughput and our results showed a robust communication (no packages were dropped) for 10 tracking sources with frequencies up to 1kHz (Figure 6). For behavioral tracking, usually two tracking sources are used, with a sampling frequency around 50 Hz. In addition, to ensure that no data are dropped, we recommend using the camera shutter events for synchronization: if the count of the shutter TTL events and the recorded tracking samples correspond, then all data have been correctly transmitted.

The `Tracking Stimulator` module also presents some limitations that will be addressed in future updates. First, the user is limited in selecting stimulation regions of circular shape, but we plan to extend to polygons and ellipses. Second, only one TTL output channel can be selected at the moment, preventing the user from triggering different stimulation regimes for different regions. In future releases, we will permit pairing of single regions with TTL outputs, which will allow the possibility, for example, of stimulating two place cells at different frequencies. However, this functionality can be achieved with the current version by using multiple `Tracking Stimulator` nodes at the same time.

In conclusion, we have presented an affordable solution for animal tracking and closed-loop stimulation using open source tools that could contribute to interrogation of neural activity, and the closed-loop investigation of complex behaviors.

## Acknowledgments

A.P.B. (PhD fellow), M.F. (PI), and P.H. (PI) are part of the Simula-UCSD-University of Oslo Research and PhD training (SUURPh) program, an international collaboration in computational biology and medicine funded by the Norwegian Ministry of Education and Research. In addition, the authors acknowledge support by the Research Council of Norway through grants no 231248 (to T.H.) and 204939, 217920, and 250259 (to M.F.).

## References

[1] Paulo Aguiar, Luís Mendonça, and Vasco Galhardo. Opencontrol: a free opensource software for video tracking and automated control of behavioral mazes. Journal of neuroscience methods, 166(1):66–72, 2007.

[2] Christopher Black, Jakob Voigts, Uday Agrawal, Max Ladow, Juan Santoyo, Christopher Moore, and Stephanie Jones. Open ephys electroencephalography (open ephys+ eeg): a modular, low-cost, open-source solution to human neural recording. Journal of Neural Engineering, 14(3):035002, 2017.

[3] György Buzsáki. Theta oscillations in the hippocampus. Neuron, 33(3):325–340, 2002.

[4] György Buzsáki. Large-scale recording of neuronal ensembles. Nature neuroscience, 7(5):446, 2004.

[5] Alison Callahan, Kim D Anderson, Michael S Beattie, John L Bixby, Adam R Ferguson, Karim Fouad, Lyn B Jakeman, Jessica L Nielson, Phillip G Popovich, Jan M Schwab, et al. Developing a data sharing community for spinal cord injury research. Experimental neurology, 295:135–143, 2017.

[6] Tsai-Wen Chen, Trevor J Wardill, Yi Sun, Stefan R Pulver, Sabine L Renninger, Amy Baohan, Eric R Schreiter, Rex A Kerr, Michael B Orger, Vivek Jayaraman, et al. Ultrasensitive fluorescent proteins for imaging neuronal activity. Nature, 499(7458):295–300, 2013.

[7] Davide Ciliberti and Fabian Kloosterman. Falcon: a highly flexible open-source software for closed-loop neuroscience. Journal of Neural Engineering, 2017.

[8] Gaetan De Lavilléon, Marie Masako Lacroix, Laure Rondi-Reig, and Karim Benchenane. Explicit memory creation during sleep demonstrates a causal role of place cells in navigation. Nature neuroscience, 18(4):493, 2015.

[9] JoséM R Delgado, Victor S Johnston, Jan D Wallace, and Ronald J Bradley. Operant conditioning of amygdala spindling in the free chimpanzee. Brain research, 22(3):347–362, 1970.

[10] Maria Diamantaki, Stefano Coletta, Khaled Nasr, Roxana Zeraati, Sophie Laturnus, Philipp Berens, Patricia Preston-Ferrer, and Andrea Burgalossi. Manipulating hippocampal place cell activity by single-cell stimulation in freely moving mice. Cell reports, 23(1):32–38, 2018.

[11] Daniel E Feldman. The spike-timing dependence of plasticity. Neuron, 75(4):556–571, 2012.

[12] Eberhard E Fetz and Dom V Finocchio. Operant conditioning of specific patterns of neural and muscular activity. Science, 174(4007):431–435, 1971.

[13] Marianne Fyhn, Sturla Molden, Stig Hollup, May-Britt Moser, and Edvard I Moser. Hippocampal neurons responding to first-time dislocation of a target object. Neuron, 35(3):555–566, 2002.

[14] Marianne Fyhn, Sturla Molden, Menno P Witter, Edvard I Moser, and May-Britt Moser. Spatial representation in the entorhinal cortex. Science, 305(5688):1258–1264, 2004.

[15] Padraig Gleeson, Andrew P Davison, R Angus Silver, and Giorgio A Ascoli. A commitment to open source in neuroscience. Neuron, 96(5):964–965, 2017.

[16] Torkel Hafting, Marianne Fyhn, Sturla Molden, May-Britt Moser, and Edvard I Moser. Microstructure of a spatial map in the entorhinal cortex. Nature, 436(7052):801–806, 2005.

[17] Brett M Hewitt, Moi Hoon Yap, Emma F Hodson-Tole, Aneurin J Kennerley, Paul S Sharp, and Robyn A Grant. A novel automated rodent tracker (art), demonstrated in a mouse model of amyotrophic lateral sclerosis. Journal of neuroscience methods, 2017.

[18] Andrew Jackson, Jaideep Mavoori, and Eberhard E Fetz. Long-term motor cortex plasticity induced by an electronic neural implant. Nature, 444(7115):56, 2006.

[19] Nathan C Klapoetke, Yasunobu Murata, Sung Soo Kim, Stefan R Pulver, Amanda Birdsey-Benson, Yong Ku Cho, Tania K Morimoto, Amy S Chuong, Eric J Carpenter, Zhijian Tian, et al. Independent optical excitation of distinct neural populations. Nature methods, 11(3):338, 2014.

[20] Emilio Kropff, James E Carmichael, May-Britt Moser, and Edvard I Moser. Speed cells in the medial entorhinal cortex. Nature, 523(7561):419–424, 2015.

[21] Joseph LeDoux and Nathaniel D Daw. Surviving threats: neural circuit and computational implications of a new taxonomy of defensive behaviour. Nature Reviews Neuroscience, 2018.

[22] Doyun Lee, Bei-Jung Lin, and Albert K Lee. Hippocampal place fields emerge upon single-cell manipulation of excitability during behavior. Science, 337(6096):849–853, 2012.

[23] Kristian Kinden Lensjø, Mikkel Elle Lepperød, Gunnar Dick, Torkel Hafting, and Marianne Fyhn. Removal of perineuronal nets unlocks juvenile plasticity through network mechanisms of decreased inhibition and increased gamma activity. Journal of Neuroscience, 37(5):1269–1283, 2017.

[24] William A Liberti III, L Nathan Perkins, Daniel P Leman, and Timothy J Gardner. An open source, wireless capable miniature microscope system. Journal of Neural Engineering, 14(4):045001, 2017.

[25] Gonçalo Lopes, Niccolò Bonacchi, João Frazão, Joana P Neto, Bassam V Atallah, Sofia Soares, Luís Moreira, Sara Matias, Pavel M Itskov, Patrícia A Correia, et al. Bonsai: an event-based framework for processing and controlling data streams. Frontiers in neuroinformatics, 9, 2015.

[26] Chet T Moritz, Steve I Perlmutter, and Eberhard E Fetz. Direct control of paralysed muscles by cortical neurons. Nature, 456(7222):639, 2008.

[27] John O’Keefe and Jonathan Dostrovsky. The hippocampus as a spatial map. preliminary evidence from unit activity in the freely-moving rat. Brain research, 34(1):171–175, 1971.

[28] Jean-Etienne Poirrier, Laurent Poirrier, Pierre Leprince, and Pierre Maquet. Gemvid, an open source, modular, automated activity recording system for rats using digital video. Journal of Circadian Rhythms, 4(1):10, 2006.

[29] James M Rebesco, Ian H Stevenson, Konrad Koerding, Sara A Solla, and Lee E Miller. Rewiring neural interactions by micro-stimulation. Frontiers in systems neuroscience, 4:39, 2010.

[30] Cyrille Rossant, Shabnam N Kadir, Dan FM Goodman, John Schulman, Maximilian LD Hunter, Aman B Saleem, Andres Grosmark, Mariano Belluscio, George H Denfield, Alexander S Ecker, et al. Spike sorting for large, dense electrode arrays. Nature neuroscience, 19(4):634–641, 2016.

[31] Andre L Samson, Lining Ju, Hyun Ah Kim, Shenpeng R Zhang, Jessica AA Lee, Sharelle A Sturgeon, Christopher G Sobey, Shaun P Jackson, and Simone M Schoenwaelder. Mousemove: an open source program for semi-automated analysis of movement and cognitive testing in rodents. Scientific reports, 5:16171, 2015.

[32] Francesca Sargolini, Marianne Fyhn, Torkel Hafting, Bruce L McNaughton, Menno P Witter, May-Britt Moser, and Edvard I Moser. Conjunctive representation of position, direction, and velocity in entorhinal cortex. Science, 312(5774):758–762, 2006.

[33] Joshua Handman Siegle, Aarón Cuevas López, Yogi Patel, Kirill Abramov, Shay Ohayon, and Jakob Voigts. Open ephys: An open-source, plugin-based platform for multichannel electrophysiology. Journal of Neural Engineering, 2017.

[34] Nicholas J Sofroniew and Karel Svoboda. Whisking. Current Biology, 25(4):R137–R140, 2015.

[35] Trygve Solstad, Charlotte N Boccara, Emilio Kropff, May-Britt Moser, and Edvard I Moser. Representation of geometric borders in the entorhinal cortex. Science, 322(5909):1865–1868, 2008.

[36] Michael A van der Kooij and Carmen Sandi. Social memories in rodents: methods, mechanisms and modulation by stress. Neuroscience & Biobehavioral Reviews, 36(7):1763–1772, 2012.

[37] Melis Yilmaz and Markus Meister. Rapid innate defensive responses of mice to looming visual stimuli. Current Biology, 23(20):2011–2015, 2013.

